# A novel *Flavobacterium quisquiliarum* porphyrin binding protein independently disrupts *Pseudomonas aeruginosa* biofilms

**DOI:** 10.64898/2026.04.07.716877

**Authors:** Ieva Lelenaite, Charlotte Stenkjær Fletcher, William Houppy, Claire Morley, Adrian Brown, Gary W. Black, Adam K. Malekpour, Nicola L. Brown, Warispreet Singh, Jose Munoz, Hamish C. L. Yau, Neil J. Lant, William G.T. Willats

## Abstract

Bacterial biofilms underpin chronic infection and antimicrobial resistance, notably in *Pseudomonas aeruginosa*. Here we deconvolute a commercial alginate lyase preparation from *Flavobacterium quisquiliarum* and identify a previously uncharacterised ∼21 kDa porphyrin-binding protein (*Fq*PBP). Structural, biophysical and docking analyses reveal high-affinity tetrapyrrole binding. Recombinant *Fq*PBP independently inhibits and disperses *P. aeruginosa* biofilms, implicating porphyrin sequestration and iron homeostasis in biofilm control and highlighting a potential therapeutic strategy targeting iron acquisition pathways.

## Main text

Bacterial biofilms are aggregations of cells embedded in a hydrated extracellular matrix that mediates attachment to biotic and abiotic surfaces and protects from immune system attack^1,2^. Biofilms also contribute to antibiotic resistance by creating favourable conditions facilitating gene transfer and by physically restricting access of antibiotics to bacteria^3,4^. They are implicated in around 80% of all human microbial infections and further impact diverse commercial and industrial sectors including personal, oral and home care, food and agriculture with associated costs totalling hundreds of billions of dollars annually^5^.

Biofilm-forming *Pseudomonas* species, including *Pseudomonas aeruginosa*, are of particular concern because of their role in serious diseases such as cystic fibrosis, which are resistant to standard antibiotic treatment^3,6,7^. The composition of *P. aeruginosa* biofilms varies according to strain and growth stage but typically contains proteins, extracellular DNA, polysaccharides, and water^8,9.^ The polysaccharide component typically contains alginate, Psl, and Pel that contribute to stabilising biofilms in mucoid phenotypes of certain strains. Alginate is one of the main virulence factors associated with *P. aeruginosa* biofilms^10,2.^ Consequently, alginate lyases (AL) have been proposed as biotherapeutic agents to disrupt *P. aeruginosa* biofilms^9^. AL treatment has been shown to reduce biofilm viscosity, and the inhibitory concentration of antibiotics against *P. aeruginosa* cultures is reduced when combined with AL activity^11,2.^ However, the mechanistic details of these effects and the precise role of AL activity are unclear^11^. For example, the synergy between tobramycin and AL appears to be decoupled from AL catalytic activity, raising the question of whether other components of AL preparations may be partly or wholly responsible for biofilm disruption^11^.

Much of the work on AL disruption of biofilms has relied on a commercial preparation from *Flavobacterium* sp. supplied by Sigma (product code A1603) and Nagase ChemteX Corporation (as ‘Alginate Lyase S’, and henceforth referred to as ‘Nagase AL’). This crude preparation contains multiple proteins, including two ALs of ∼30 kDa (Aly30) and ∼40 kDa (Aly40) identified by Takeuchi *et al*. and Blanco-Cabra^2,12,13^. Blanco-Cabra *et al*. also identified two proteins of ∼21 kDa and ∼46 kDa in Nagase AL but did not determine their identity or functions^2^. Morley *et al*. recently confirmed that Nagase AL is essentially a cell-free extract from *Flavobacterium quisquiliarum*^14^. Here, we further deconvoluted the constituent proteins in Nagase AL and investigated their roles in *P. aeruginosa* biofilm dispersal.

SDS-PAGE analysis of Nagase AL and Sigma A1603 yielded nearly identical bands, including ones corresponding to the previously identified ∼30 kDa and ∼40 kDa ALs and the ∼21 kDa protein band of unknown function (**Figure 1A**). The ∼21 kDa band was excised and constituent proteins identified using standard discovery proteomic methods^15^. The highest-scoring and most abundant identified protein was NCBI accession WP_179002276.1, from *F. quisquiliarum* (**Supplemental Table 1**). Sequence-based searches did not yield homologs of WP_179002276.1 or suggest a function, so we instead searched for structurally similar proteins. The AlphaFold2 structure of WP_179002276.1 was submitted as a PDB file to the Dali server for comparison with 3D structures deposited in the Protein Data Bank^16^. The top-ranked match was the HusA hemophore from *Porphyromonas gingivalis* (PDB ID 6BQS). WP_179002276.1 has only limited amino acid sequence identity (24.3%) with HusA (**Supplemental Figure 1**). However, AlphaFold2 prediction modelling of WP_179002276.1 compared to an NMR-derived structure of HusA confirmed a high degree of structural similarity (**Figure 1B**) with a root mean square deviation (alpha-carbons) of 1.558 Å. WP_179002276.1 will henceforth be referred to as *Fq*PBP, and we hypothesised that *Fq*PBP may fulfil a similar role to the well-documented porphyrin-binding, heme-acquisition role of HusA.

**Figure 1.**
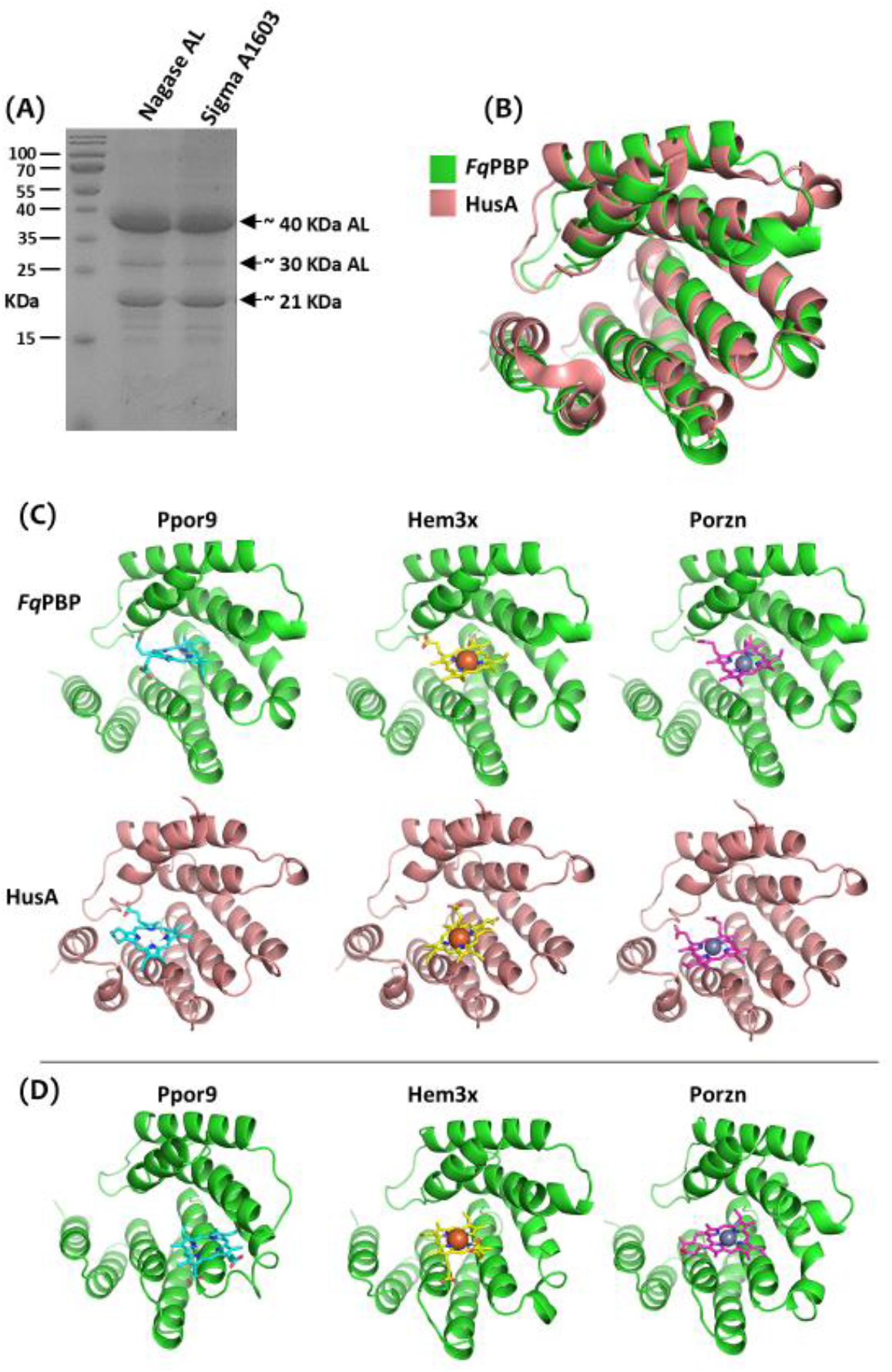
**(A)** SDS-PAGE gel showing the constituent proteins in crude alginate lyase preparations from *Flavobacterium* sp. supplied by Nagase (Nagase AL) and Sigma (A1603). In addition to the bands corresponding to ∼40 kDa and ∼30 kDa alginate lyases (ALs), note the prominent band at ∼21 kDa, subsequently identified by proteomics as WP_179002276.1 from *Flavobacterium quisquiliarum* (*Fq*PBP). **(B)** AlphaFold2-predicted structure of *Fq*PBP (green) superimposed on the NMR-derived structure of HusA (beige) showing the high structural similarity. **(C)** Molecular docking of protoporphyrin IX (Ppor9), hemin (Hem3x), and zinc protoporphyrin (Porzn) ligands to the 3D structures of *Fq*PBP and HusA. **(D)** The most populated structures obtained by cluster analysis of the equilibrated trajectory of *Fq*PBP in complex with protoporphyrin IX, hemin, and zinc protoporphyrin.

Molecular docking revealed a clear preference of *Fq*PBP for tetrapyrrole macrocycles. Protoporphyrin IX zinc complex (−10.5 kcal/mol) and protoporphyrin IX (−10.5 kcal/mol) exhibited the most favourable binding energies, followed by hemin (−10.2 kcal/mol) and heme B (−10.2 kcal/mol) (**Supplemental Table 2** and **Figure 1C**). Comparative docking demonstrated that *Fq*PBP consistently achieved stronger binding scores than HusA (−7.9 to −8.5 kcal/mol for equivalent ligands), suggesting enhanced porphyrin affinity (**Supplemental Table 2** and **Figure 1C**). All three ligands adopted similar binding orientations within a hydrophobic groove formed between α-helices, with the porphyrin macrocycle positioned to facilitate π-stacking interactions (**Figure 1C**). The biosynthetic precursors coproporphyrinogen III (−8.5 kcal/mol) and protoporphyrinogen IX (−9.0 kcal/mol) displayed weaker binding, reflecting *Fq*PBP’s preference for fully conjugated, planar aromatic systems that enable enhanced π-stacking within the binding pocket (**Supplemental Table 2)**. Molecular dynamics simulations (4 × 250 ns per complex) confirmed stable ligand binding, with backbone root mean square deviation (RMSD) values plateauing at 1–3 Å across all replicates. Cluster analysis (**Figure 1D**) revealed consistent binding poses throughout the trajectories for protoporphyrin IX, hemin, and zinc protoporphyrin. RMSF analysis demonstrated that porphyrin binding substantially reduces protein flexibility, with fluctuations decreasing by 2– 4 Å in loop regions (residues 40–60) and the C-terminal region (beyond residue 170) compared to apo-*Fq*PBP (**Figure 1D**). This ligand-induced rigidification was reproducible across independent replicates, suggesting that porphyrin binding stabilises *Fq*PBP through restriction of conformational dynamics in key flexible regions.

Molecular Mechanics-Poisson-Boltzmann Surface Area (MM-PBSA) calculations with protoporphyrin IX identified key residues contributing to binding: Arg179 (−4.64 kcal/mol) and Lys142 (−2.69 kcal/mol) provide electrostatic anchoring to porphyrin carboxylates, while Phe53 and Trp143 engage in π-stacking interactions with the conjugated tetrapyrrole macrocycle^17^ and Val97, Val98, Pro101, and Gly139 contribute additional hydrophobic contacts (**Figure 2A and 2B**). Notably, Asp135 exhibited a particularly unfavourable contribution (+2.80 kcal/mol), likely due to electrostatic repulsion with porphyrin carboxylates along with Glu178, while enabling a structure-stabilising salt bridge with Arg179 between two helices. This binding architecture, combining electrostatic anchoring through basic residues with hydrophobic stabilisation and π-stacking of the macrocycle, resembles mechanisms described for the structurally related HusA hemophore from *Porphyromonas gingivalis* (RMSD 1.558 Å between *Fq*PBP and HusA structures)^18^. Computational predictions of *Fq*PBP porphyrin binding were confirmed empirically using UV/Vis spectroscopy with recombinant *Fq*PBP, which showed characteristic spectral shifts upon binding protoporphyrin IX, zinc protoporphyrin, and hemin at 1:1 molar ratios (**Figure 2C**). Taken together, the *in silico* and empirical data provide evidence that *Fq*PBP, like HusA, binds both metalated and metal-free porphyrins without direct iron coordination, supporting a potential role in porphyrin acquisition pathways.

**Figure 2.**
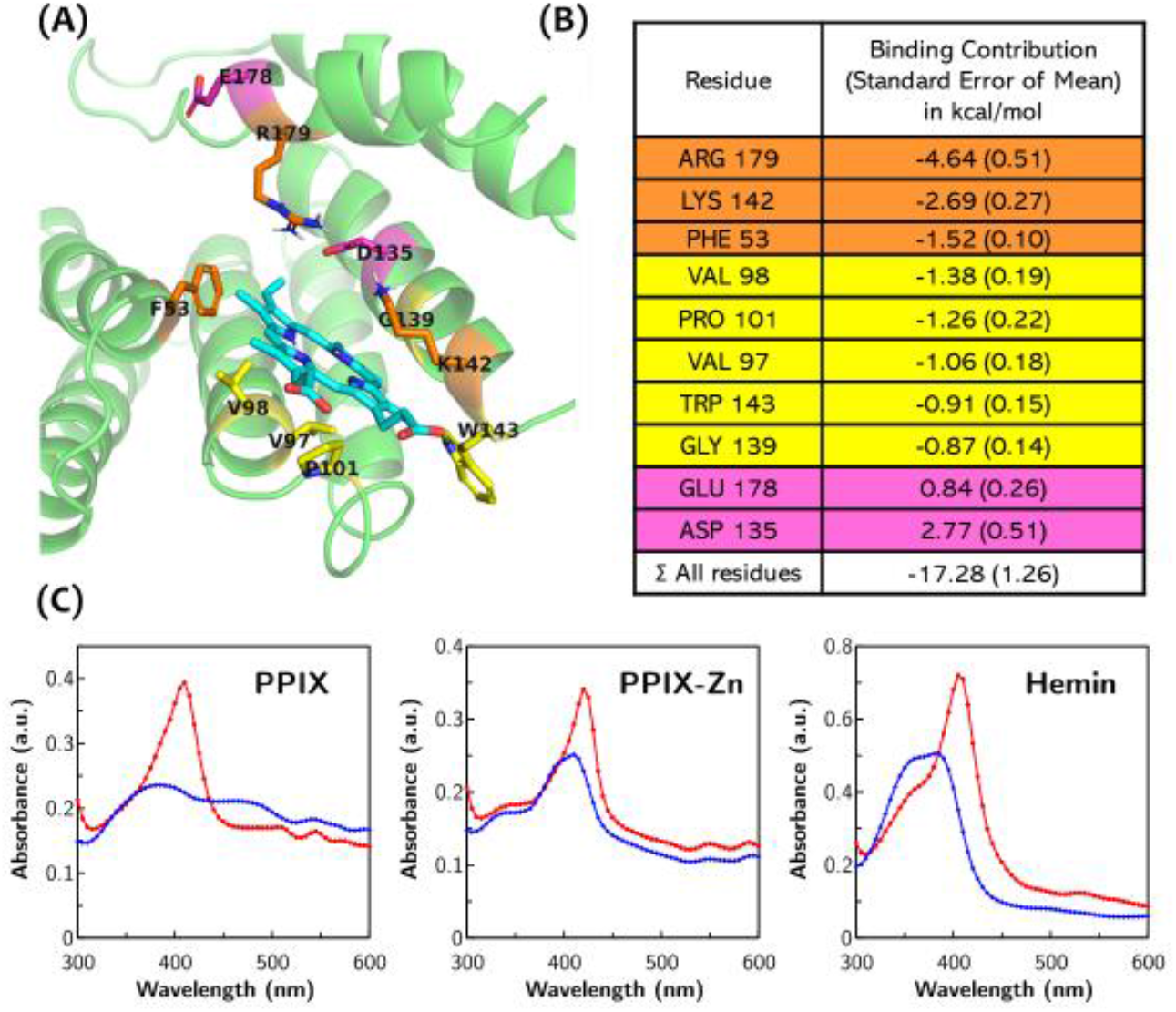
**(A)** Cluster model obtained from MD simulations of *Fq*PBP in complex with protoporphyrin IX. Important binding-site residues are highlighted in liquorice representation and colour-coded according to per-residue energy contributions obtained from MM-PBSA calculations. **(B)** Molecular Mechanics Poisson–Boltzmann Surface Area (MM-PBSA) calculations to estimate the binding free energy and decompose it into major per-residue contributions for residues in the binding site of *Fq*PBP in complex with protoporphyrin IX, obtained from four 250 ns trajectories. **(C)** UV/Vis spectroscopy confirming the porphyrin-binding capacity of recombinant *Fq*PBP against protoporphyrin IX (PPIX), its Zn complex (PPIX–Zn), and hemin at a 1:1 molar ratio. In all cases, the spectra of these porphyrins show the expected change in UV/Vis spectrum (blue to red) on addition of the protein.

The capacity of recombinant *Fq*PBP to disperse established biofilms was tested using crystal violet assays with *P. aeruginosa* PA19882 as the biofilm-producing organism. Both Nagase AL and *Fq*PBP alone dispersed established biofilms and inhibited the formation of new biofilms (**Figure 3A**). Biofilm dispersal by *Fq*PBP was greatly reduced by boiling (**Figure 3B**). Biofilm dispersal activity of *Fq*PBP was tested in combination with various permutations of recombinant versions of AL30 and AL40 (**Figure 3C**). *Fq*PBP biofilm dispersal activity was not significantly altered when used in combination with AL30 alone, AL40 alone, or AL30 and AL40 together (**Figure 3C**), confirming that *Fq*PBP is capable of biofilm dispersal independently of, and at a similar level to, AL alone. Dot blots of biofilm extracts confirmed that both alginate and a ligand recognised by *Fq*PBP are present in *P. aeruginosa* biofilms (**Figure 3D**).

**Figure 3.**
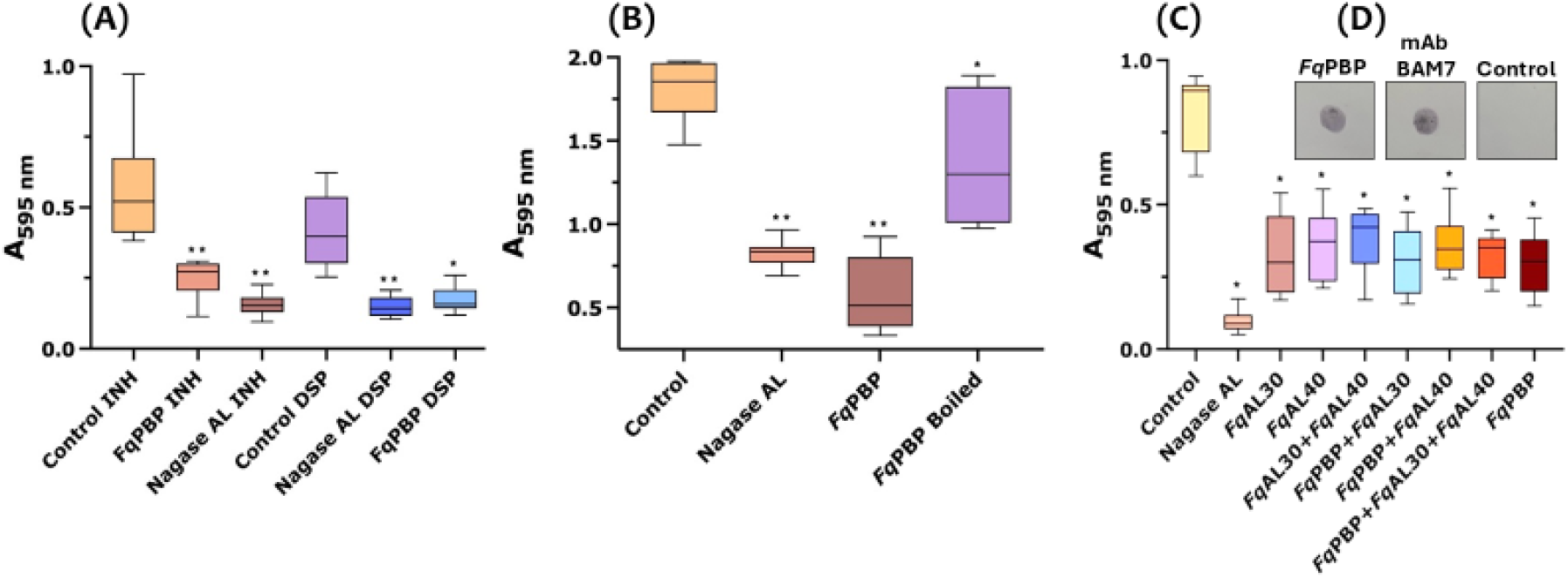
Biofilm dispersal assays using crystal violet detection of *P. aeruginosa* biofilms. **(A)** The capacity of *Fq*PBP and Nagase AL to inhibit biofilm formation (INH) and to disperse mature biofilms (DSP) was tested. **(B)** The biofilm dispersal activity of *Fq*PBP was greatly reduced by boiling. **(C)** Biofilm dispersal activity of *Fq*PBP was similar to that of the recombinant alginate lyases *Fq*AL30 and *Fq*AL40 and to combinations of *Fq*AL30 and *Fq*AL40 with *Fq*PBP. **(D)** Dot blots of material extracted from *P. aeruginosa* biofilm confirmed the presence of a *Fq*PBP ligand (detected using His-tagged *Fq*PBP) and alginate (detected using the anti-alginate antibody BAM7). In all cases, biofilms were treated with proteins at 37 °C and 100 rpm for 3 h. Each treatment consisted of 8 replicates. *Fq*PBP was used at 5%, and Nagase AL was used at 12.5 mg/mL. Unpaired t-tests were performed to compare the means of untreated control and treated biofilms; * p < 0.05, ** p < 0.0001. Control treatments were buffer only.

Taken together, these data support the hypothesis that *Fq*PBP is a novel PBP with biofilm dispersal activity. The fact that *Fq*PBP both inhibits *P. aeruginosa* biofilm formation and disperses established biofilms suggests it may interfere with regulatory aspects of biofilm production and directly or indirectly impact biofilm structural integrity post-synthesis via porphyrin binding. Iron is required for *P. aeruginosa* biofilm formation, and since iron chelators disrupt *P. aeruginosa* biofilms^19,20,21,22,23^, iron sequestration by *Fq*PBP may similarly inhibit biofilm formation. *Pseudomonas* spp. and *Flavobacterium* spp. frequently occur in mixed-species biofilms in which they compete for available resources^24^. Like HusA, *Fq*PBP may act as a heme scavenger, helping to secure iron within ecological niches where it may be scarce, but more work is required to better understand *Fq*PBP biology. Developing more effective strategies for tackling biofilms is a medical priority, and the therapeutic potential of targeting iron acquisition is demonstrated by iron-chelator medicines including Deferasirox and Deferoxamine^25^. Since porphyrins are components of biofilms produced by a wide range of microbes, dispersal mechanisms based on porphyrin binding may have broader applications beyond *Pseudomonas* spp.

This work is also significant in relation to developing a better understanding of *Flavobacterium* spp. virulence. Several *Flavobacterium* spp., and especially *Flavobacterium columnare*, which causes columnaris disease, are economically devastating for the fish-farming industry^26,27.^ Current control methods are inadequate, and virulence mechanisms of *F. columnare* are not well understood. Iron acquisition is required for *F. columnare* infection, and multiple components of the *F. columnare* iron uptake systems have been identified^26^. However, *F. columnare* mutants that failed to produce siderophores and other iron acquisition functions remain virulent^28^. It is possible that redundancies among the iron acquisition systems are responsible for these effects, but the identification of a novel *Flavobacterium* sp. PBP provides insight into a potentially additional component of *F. columnare* iron uptake and virulence.

## Methods

### In silico structural analysis

The three-dimensional structure of the *F. quisquiliarum* porphyrin-binding protein, *Fq*PBP (WP_179002276.1), was predicted using AlphaFold2 via the ColabFold server, while the 3D structure of HusA was retrieved from the PDB database (PDB ID: 6BQS)^29,30.^ The protonation states of titratable amino acid residues were determined at pH 7.0 using the H++ server^31^. The structures of protoporphyrin IX, zinc protoporphyrin, and hemin were retrieved from the PubChem database and prepared for docking using AutoDockTools^32^. Gasteiger charges were assigned to all ligands, and geometry optimisation was performed prior to docking.

### In silico molecular docking

Molecular docking was performed using AutoDock Vina^33^. The grid box was centred on the putative binding site of *Fq*PBP and HusA with dimensions of 23.25 × 21.0 × 24.75 Å. The exhaustiveness parameter was set to 32 to ensure thorough sampling of conformational space. Nine binding poses were generated for each ligand, and poses were ranked according to their docking scores (kcal/mol). The docking results were visualised and analysed using PyMOL (Schrödinger, LLC).

### In silico molecular dynamics simulations

Molecular dynamics (MD) simulations were performed using the GPU-accelerated version of PMEMD implemented in Amber 20. The ff19SB force field was used for the protein, and the General Amber Force Field (GAFF) was employed for the porphyrin ligands^34,35.^ For hemin and zinc protoporphyrin, force-field parameters were generated using the MCPB.py module, with geometry optimisation and charge calculations performed at the B3LYP/6-31G(d) level of theory using Gaussian 16^36^. Partial charges for protoporphyrin IX were calculated using the restrained electrostatic potential (RESP) method. Each complex was solvated in a truncated cubic box of TIP3P water molecules extending 10 Å from the protein surface. Counterions were added to neutralise the system.

#### The simulation protocol consisted of the following stages

##### Energy minimization

The system was subjected to energy minimisation for 1,000 cycles, with the first 500 cycles using the steepest descent method followed by conjugate gradient minimisation. Harmonic restraints of 2.0 kcal/mol/Å^2^ were applied to all protein residues during minimisation. Heating: The system was gradually heated from 0 K to 298.15 K over 50 ps under constant volume (NVT ensemble) using a Langevin thermostat with a collision frequency of 2.0 ps^−1^. Restraints of 2.0 kcal/mol/Å^2^ were maintained on all protein residues.

##### Density Equilibration

Pressure equilibration was performed for 50 ps under constant pressure (NPT ensemble) at 1 bar using isotropic position scaling with a pressure relaxation time of 1.0 ps. Restraints of 2.0 kcal/mol/Å^2^ were maintained on all protein residues.

##### Equilibration

Further equilibration was conducted for 500 ps under the NPT ensemble at 298.15 K and 1 bar, with a pressure relaxation time of 2.0 ps, without positional restraints.

##### Production MD

Four independent replica simulations of 250 ns each were performed for each system, providing a cumulative sampling of 1 μs. Production runs were conducted under the NPT ensemble at 298.15 K and 1 bar using a Berendsen barostat with a pressure relaxation time of 0.5 ps. The SHAKE algorithm was applied to constrain bonds involving hydrogen atoms, allowing a time step of 2 fs. Long-range electrostatic interactions were calculated using the particle mesh Ewald (PME) method with a non-bonded cutoff of 12 Å. Trajectory coordinates were saved every 10 ps (5,000 steps) for subsequent analysis. The stability of the protein and protein–ligand complexes was assessed by calculating the root mean square deviation (RMSD) of backbone atoms relative to the initial structure. Root mean square fluctuation (RMSF) analysis was performed to identify flexible regions within the protein. Cluster analysis was conducted to identify representative conformations from the MD trajectories. All analyses were performed using the cpptraj module of AmberTools. Molecular dynamics trajectories were visualised using Visual Molecular Dynamics.

### Binding Free Energy Calculations

Binding free energies were calculated using the Molecular Mechanics Poisson–Boltzmann Surface Area (MM-PBSA) method implemented in the MMPBSA.py module of AmberTools. A total of 2,000 frames were extracted from the equilibrated portion of the MD trajectories (50–250 ns) at intervals of every 10 frames for analysis. The binding free energy (ΔG_bind) was decomposed into contributions from van der Waals interactions, electrostatic interactions, polar solvation energy, and non-polar solvation energy. Per-residue energy decomposition (idecomp=1) was performed to identify key residues contributing to ligand binding.

### UV/Vis spectroscopy

Protein solutions at a concentration of 15 μM were prepared in 50 mM Tris-HCl (Sigma Aldrich UK, T5941) with 150 mM NaCl (VWR UK, 27810.295) buffer in de-ionised water. Porphyrin substrates (protoporphyrin IX (Sigma Aldrich UK, 258385), protoporphyrin IX-Zn (Sigma Aldrich UK, 282820), hemin (Sigma Aldrich, H9039)) were prepared at a concentration of 15 μM in 0.1 M NaOH. One millilitre of protein solution was combined with 1 mL of porphyrin substrate in a separate 2 mL Eppendorf tube and mixed by hand. For controls without protein, 1 mL of buffer solution was used. From these solutions, 200 μL aliquots were taken and transferred to a clear 96-well plate, which was then incubated at 25 °C for 10 min. A spectrum scan was then performed, measuring the absorbance from 300 nm to 600 nm, using a Molecular Devices Spectramax M2e microplate reader.

### LC-MS analysis of Nagase gel bands

Protein samples were mixed with LDS sample buffer (Thermo Scientific) and resolved on a 5–20% polyacrylamide gradient gel (mPAGE, Merck) using Tris-MOPS running buffer. Proteins were stained with colloidal Coomassie, and protein bands were excised for in-gel trypsin or chymotrypsin digestion. This was performed with sequencing-grade enzymes and ProteaseMax surfactant trypsin enhancer (Promega V5111, V1061, and V2071) using the manufacturers’ protocols. Recovered peptides were desalted using C18 StageTips and LC-MS analysis was carried out in trap-and-elute mode on a Sciex TripleTOF 6600 linked to an Eksigent 425 nanoLC system via a Duospray II source^15^. A solvent flow of 5 μL/min was used over YMC TriArt C18 columns (resolving: 1/32”, S-3 mm, 150 × 0.3 mm, TA12S03-15H0RU; trap: 1/32”, 5 mm, 5 × 0.5 mm, TA12S05-E5J0RU). Top-10 data-dependent acquisition, with a cycle time of 1.3 s, was performed for a total of 23.2 min during gradients of 5–35% B over 20 min and 35–80% B over 2 min before a 4-min hold at 80% B (solvent A: 0.1% formic acid in H2O; solvent B: 0.1% formic acid in acetonitrile).

Sciex wiff-format acquired data files were processed with PEAKS Studio 10.6 software (Bioinformatics Solutions Inc.). Settings for peptide and protein identification were: semi-specific trypsin digestion; fixed residue modification, carbamidomethyl (cysteine); and variable modification, oxidised (methionine). Other parameters were default software values assigned for a TripleTOF instrument. The search database comprised 68,733 entries and contained the *Arabidopsis thaliana* proteome, known proteomic experiment contaminants, bacterial proteomes from *Flavobacterium collinsii, Flavobacterium phragmatis, Flavobacterium crocinum, Zobellia galactanivorans* and *Sphingobacterium multivorum*, and a combined *Flavobacterium* protein download from NCBI (December 2021; 4,540 sequences). Post-processing, the peptide:spectrum match FDR was set at 0.1%, the ALC score threshold was set to >80%, and a filter of two significant peptides per protein was applied.

### Biofilm dispersal assays

To examine biofilm dispersal *in vitro*, a crystal violet (CV) microtitre assay was developed. Briefly, *Pseudomonas aeruginosa* (DSMZ 19882) was grown overnight in Luria-Bertani (Miller) at 37 °C with agitation (150 rpm). The overnight culture was diluted 1:100 (v/v) in LB and an aliquot of 200 μL was transferred to a polystyrene 96-well microtitre plate. Biofilms were allowed to establish at 37 °C under static conditions for 26 h. Negative controls consisted of non-inoculated medium, incubated under the same conditions. Established biofilms were treated by adding dispersal agents at 5% (10 μL of agent to 200 μL culture) and incubating at 37 °C with agitation (100 rpm) for 3 h. Following the incubation period, spent media and unbound planktonic cells were removed by inversion and rinsed thoroughly with MilliQ (MQ) water. Residual bacterial material and adherent biofilms were stained with 0.1% Crystal Violet (CV) (250 μL) for 15 min. Excess and unbound dye was thoroughly washed away with MQ water. Bound dye was solubilised with ethanol (97%) and acetic acid (3%) solution (250 μL), and biofilm formation/dispersal was determined spectrophotometrically (A595 nm) with a plate reader. Biofilm formation/dispersal assays were carried out with eight replicates per treatment. Absorbance values were background-corrected by subtracting the absorbance values of the negative-control wells. Statistical analysis involved performing pairwise t-tests to compare the means of untreated control and treated biofilms. Graphs and statistical analyses were performed in GraphPad Prism.

### Production of recombinant Nagase AL constituent proteins

Plasmids encoding PBPs were transformed into *E. coli* BL21(DE3) chemically competent cells and grown in LB broth at 37 °C supplemented with 50 μg/mL kanamycin. Production of recombinant proteins was induced by the addition of 0.5 mM IPTG at a culture OD600 = 0.6, and incubation at 16 °C for 16 h. To purify PBPs, cells were collected by centrifugation, resuspended in 1× Talon buffer (20 mM Tris, 300 mM NaCl, pH 8.0), and lysed by sonication. The resulting cell-free extracts were then centrifuged for 30 min at 14,600 rpm and purified via immobilised metal affinity chromatography (IMAC) using a cobalt-based metal affinity matrix (Talon IMAC resin) by stepwise elution with imidazole. Positive fractions, identified by SDS-PAGE gel electrophoresis, were dialysed into 20 mM Tris base, 100 mM NaCl, pH 7.5 and concentrated using a stirred-cell Amicon with a 10 kDa cut-off.

### Dot blots of Pseudomonas aeruginosa extracts

*Pseudomonas aeruginosa* (DSMZ 19882) biofilms were either spotted directly onto the nitrocellulose membrane or scraped and reconstituted with ddH2O, 50 mM CDTA (pH 7.5), and 4 M NaOH + 0.1% NaBH4 (w/v), then sonicated for 10 min in an ultrasonic bath prior to spotting onto the nitrocellulose membrane. The membranes were then blocked with 5% milk in PBS and probed with either BAM7 or WP_179002276.1. Dot blots were then probed with secondary antibodies (anti-rat for BAM7 or anti-polyhistidine for WP_179002276.1) and developed using NBT/BCIP colour development solution.

## Author contributions

**IL**: Produced recombinant proteins; designed, performed, and analysed biofilm dispersal assays; and contributed to writing the manuscript and preparing figures.**CF**: Produced recombinant proteins; designed, performed, and analysed biofilm dispersal assays; performed dot blots; and contributed to writing the manuscript and preparing figures.**WH**: Performed computational protein analysis; contributed to designing experiments; analysed data; and contributed to writing the manuscript and preparing figures.**CM**: Performed SDS-PAGE gel, contributed to designing experiments and reviewing the manuscript.

**AB**: Designed and performed proteomics experiments.

**GB**: Designed experiments; analysed data; and reviewed the manuscript.**AM**: Designed, performed and analysed porphyrin binding tests.

**NB**: Cloned expression constructs, produced recombinant proteins for testing, reviewed manuscript.

**WS**: Performed computational protein analysis; contributed to designing experiments; analysed data; and contributed to writing the manuscript and preparing figures.**JM:** Contributed to designing experiments and reviewing the manuscript.

**HY:** Contributed to designing experiments; analysed data; and contributed to writing the manuscript and preparing figures.**NL**: Contributed to designing experiments; performed computational protein analysis; analysed data; and contributed to writing the manuscript and preparing figures. **WW**: Contributed to designing experiments; analysed data; and contributed to writing the manuscript and preparing figures.

## Acknowledgements

We thank Adam Azzi and Ewelina Bien for experimental contributions, and the BBSRC for funding in relation to the BiSCoP (Bioscience for Sustainable Consumer Products) Collaborative Training Partnership.

## Figure legends

**Supplemental Table 1**. Identity of constituent proteins within the ∼21 kDa band separated from Nagase crude alginate lyase preparation.

**Supplemental Table 2**. Docking scores (kcal/mol) of *Fq*PBP and HusA against porphyrin related ligands obtained from AutoDock Vina (More negative values indicate higher binding affinity).

**Supplemental Figure 1**. Sequence alignment of *Fq*PBP and HusA generated with ClustalO including the secondary structures. The major binding contributing residues found by MMPBSA with Protoporphyrin IX on *Fq*PBP are shown below the sequence by the symbols 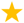, 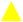 and 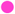. The marks ***1*** indicate a cysteine bridge in *Fq*PBP.

## References

1. Haidar, A., Muazzam, A., Nadeem, A., Atique, R., Saeed, H.A., Naveed, A., Sharif, J., Perveen, A., Fatima, H.R. and Samad, A. Biofilm formation and antibiotic resistance in Pseudomonas aeruginosa. The Microbe 3, 100078 (2024).

2. Blanco-Cabra, N., Paetzold, B., Ferrar, T., Mazzolini, R., Torrents, E., Serrano, L. and Lluch-Senar, M. Characterization of different alginate lyases for dissolving Pseudomonas aeruginosa biofilms. Scientific Reports 10(1), 9390 (2020).

3. Vetrivel, A., Ramasamy, M., Vetrivel, P., Natchimuthu, S., Arunachalam, S., Kim, G.S. and Murugesan, R. Pseudomonas aeruginosa biofilm formation and its control. Biologics 1(3), 312–336 (2021).

4. Høiby, N., Bjarnsholt, T., Givskov, M., Molin, S. and Ciofu, O. Antibiotic resistance of bacterial biofilms. International journal of antimicrobial agents 35(4), 322–332 (2010).

5. Cámara, M., Green, W., MacPhee, C.E., Rakowska, P.D., Raval, R., Richardson, M.C., Slater-Jefferies, J., Steventon, K. and Webb, J.S. Economic significance of biofilms: a multidisciplinary and cross-sectoral challenge. npj Biofilms and Microbiomes 8(1), 42 (2022).

6. Stover, C.K., Pham, X.Q., Erwin, A.L., Mizoguchi, S.D., Warrener, P., Hickey, M.J., Brinkman, F.S.L., Hufnagle, W.O., Kowalik, D.J., Lagrou, M. and Garber, R.L. Complete genome sequence of Pseudomonas aeruginosa PAO1, an opportunistic pathogen. Nature 406(6799), 959–964 (2000).

7. Tuon, F.F., Dantas, L.R., Suss, P.H. and Tasca Ribeiro, V.S. Pathogenesis of the Pseudomonas aeruginosa biofilm: a review. Pathogens 11(3), 300 (2022).

8. Mann, E.E. and Wozniak, D.J. Pseudomonas biofilm matrix composition and niche biology. FEMS microbiology reviews 36(4), 893–916 (2012).

9. Wang, S., Zhao, Y., Breslawec, A.P., Liang, T., Deng, Z., Kuperman, L.L. and Yu, Q. Strategy to combat biofilms: a focus on biofilm dispersal enzymes. npj Biofilms and Microbiomes 9(1), 63 (2023).

10. Boyd, A. and Chakrabarty, A.M. Pseudomonas aeruginosa biofilms: role of the alginate exopolysaccharide. Journal of industrial microbiology and biotechnology 15(3), 162–168 (1995).

11. Lamppa, J.W. and Griswold, K.E. Alginate lyase exhibits catalysis-independent biofilm dispersion and antibiotic synergy. Antimicrobial agents and chemotherapy 57(1), 137–145 (2013).

12. Takeuchi, T., Murata, K. and Kusakabe, I. A method for depolymerization of alginate using the enzyme system of Flavobacterium multivolum. Nippon Shokuhin Kogyo Gakkaishi 41(7), 505–511 (1994).

13. Takeuchi, T., Nibu, Y., Murata, K. and Kusakabe, I. Purification and Characterization of Endo Poly (α-L-Guluronate) Lyase in the Enzyme System from Flavobacterium multivolum. Food Science and Technology International, Tokyo 3(1), 22–26 (1997).

14. Morley, C., Yau, H.C., Houppy, W., Singh, W., Lant, N.J., Black, G.W. and Munoz-Munoz, J. Structure/activity relationships of two alginate lyases from Flavobacterium spp. and their potential application in detergents. International Journal of Biological Macromolecules 310, 143524 (2025).

15. Rappsilber, J., Mann, M. and Ishihama, Y. Protocol for micro-purification, enrichment, pre-fractionation and storage of peptides for proteomics using StageTips. Nature protocols 2(8), 1896–1906 (2007).

16. Holm, L. Dali server: structural unification of protein families. Nucleic acids research 50(W1), W210–W215 (2022).

17. Li, T., Bonkovsky, H.L. and Gua, J. Structural analysis of heme proteins: implications for design and prediction. BMC Structural Biology 11(13), 1–13 (2011)

18. Gao, J.L., Nguyen, K.A. and Hunter, N. Characterization of a hemophore-like protein from Porphyromonas gingivalis. Journal of Biological Chemistry 285(51), 40028–40038 (2010).

19. Banin, E., Vasil, M.L. and Greenberg, E.P. Iron and Pseudomonas aeruginosa biofilm formation. Proceedings of the National Academy of Sciences 102(31), 11076–11081 (2005).

20. Oliveira, F., Rohde, H., Vilanova, M. and Cerca, N. The emerging role of iron acquisition in biofilm-associated infections. Trends in microbiology 29(9), 772–775 (2021).

21. Jones, L.M., Dunham, D., Rennie, M.Y., Kirman, J., Lopez, A.J., Keim, K.C., Little, W., Gomez, A., Bourke, J., Ng, H. and DaCosta, R.S. In vitro detection of porphyrin-producing wound bacteria with real-time fluorescence imaging. Future microbiology 15(5), 319–332 (2020).

22. Singh, P.K., Parsek, M.R., Greenberg, E.P. and Welsh, M.J. A component of innate immunity prevents bacterial biofilm development. Nature 417(6888), 552–555 (2002).

23. Ganchev, I. Role of Iron Homeostasis in the Multispecies Biofilm Formation. Microbiology 92(5), 675–685 (2023).

24. Zhang, W., Sileika, T. and Packman, A.I. Effects of fluid flow conditions on interactions between species in biofilms. FEMS microbiology ecology 84(2), 344–354 (2013).

25. Moreau-Marquis, S., O’Toole, G.A. and Stanton, B.A. Tobramycin and FDA-Approved Iron Chelators Eliminate Pseudomonas aeruginosa Biofilms on Cystic Fibrosis Cells. American Journal of Respiratory Cell and Molecular Biology 41, 305–313 (2009).

26. Conrad, R.A., Evenhuis, J.P., Lipscomb, R.S., Perez-Pascual, D., Stevick, R.J., Birkett, C., Ghigo, J.M. and McBride, M.J. Flavobacterium columnare ferric iron uptake systems are required for virulence. Frontiers in Cellular and Infection Microbiology 12, 1029833 (2022).

27. Fujiwara-Nagata, E., Rochat, T., Lee, B.H., Lallias, D., Rigaudeau, D. and Duchaud, E. Host specificity and virulence of Flavobacterium psychrophilum: a comparative study in ayu (Plecoglossus altivelis) and rainbow trout (Oncorhynchus mykiss) hosts. Veterinary Research 55(1), 75 (2024).

28. Conrad, R.A., Evenhuis, J.P., Lipscomb, R.S., Birkett, C. and McBride, M.J. Siderophores produced by the fish pathogen Flavobacterium columnare strain MS-FC-4 are not essential for its virulence. Applied and Environmental Microbiology 88(17), e00948–22 (2022).

29. Mirdita, M., Schütze, K., Moriwaki, Y., Heo, L., Ovchinnikov, S. and Steinegger, M. ColabFold: making protein folding accessible to all. Nature methods 19(6), 679–682 (2022).

30. Gao, J.L., Kwan, A.H., Yammine, A., Zhou, X., Trewhella, J., Hugrass, B.M., Collins, D.A., Horne, J., Ye, P., Harty, D. and Nguyen, K.A. Structural properties of a haemophore facilitate targeted elimination of the pathogen Porphyromonas gingivalis. Nature Communications 9(1), 4097 (2018).

31. Anandakrishnan, R., Aguilar, B. and Onufriev, A.V. H++ 3.0: automating p K prediction and the preparation of biomolecular structures for atomistic molecular modeling and simulations. Nucleic acids research 40(W1), W537–W541 (2012).

32. Morris, G.M., Huey, R., Lindstrom, W., Sanner, M.F., Belew, R.K., Goodsell, D.S. and Olson, A.J.AutoDock4 and AutoDockTools4: Automated docking with selective receptor flexibility. Journal of computational chemistry 30(16), 2785–2791 (2009).

33. Trott, O. and Olson, A.J.J.J.O.C.C. Software news and update AutoDock Vina: Improving the speed and accuracy of docking with a new scoring function. Effic. Optim. Multithreading 31, 455–461 (2009).

34. Tian, C., Kasavajhala, K., Belfon, K.A., Raguette, L., Huang, H., Migues, A.N., Bickel, J., Wang, Y., Pincay, J., Wu, Q. and Simmerling, C. ff19SB: amino-acid-specific protein backbone parameters trained against quantum mechanics energy surfaces in solution. Journal of chemical theory and computation 16(1), 528–552 (2019).

35. Wang, J., Wolf, R.M., Caldwell, J.W., Kollman, P.A. and Case, D.A. Development and testing of a general amber force field. Journal of computational chemistry 25(9), 1157–1174 (2004).

36. Li, P. and Merz Jr, K.M. MCPB. py: A python based metal center parameter builder. Journal of Chemical Information and Modelling 56(4), 599–604 (2016).

