## Supplemental material for "A novel *Flavobacterium quisquiliarum* porphyrin binding protein independently disrupts *Pseudomonas aeruginosa* biofilms"

### Supplemental Table 1

| Protein Group | Accession | -10lgP | Coverage (%) | Area Sum | #Peptides | #Unique | Avg. Mass | Description |
|---|---|---|---|---|---|---|---|---|
| 1 | WP_179002276.1 | 386.98 | 88 | 1.17E+07 | 133 | 46 | 22828 | MULTISPECIES: hypothetic |
| 3 | WP_179006134.1 | 321.45 | 78 | 2.20E+06 | 67 | 67 | 48420 | polysaccharide lyase famil |
| 5 | cont82 | 224.23 | 50 | 3.79E+06 | 31 | 26 | 25666 | gi117615 \| sp P00766 CTR |
| 7 | NWL03306.1 | 216.91 | 62 | 6.53E+04 | 20 | 20 | 32910 | polysaccharide lyase famil |
| 7 | WP_179006140.1 | 216.91 | 62 | 6.53E+04 | 20 | 20 | 32910 | MULTISPECIES: polysaccha |
| 6 | cont143 | 208.43 | 36 | 2.74E+06 | 16 | 16 | 23474 | pdb1FNI_A A Chain A Crys |
| 6 | cont40 | 208.43 | 36 | 2.74E+06 | 16 | 16 | 23476 | gi2914482 \| pdb 1TFX A Ch |
| 6 | cont23 | 208.43 | 36 | 2.74E+06 | 16 | 16 | 23474 | gi494360 \| pdb 1MCT A Ch |
| 6 | cont141 | 208.43 | 35 | 2.74E+06 | 16 | 16 | 24409 | sptP00761 Trypsin precurs |
| 6 | P00761 | 208.43 | 35 | 2.74E+06 | 16 | 16 | 24409 | SWISS-PROT:P00761\|TRY |
| 9 | WP_179002398.1 | 187.68 | 69 | 8.06E+04 | 20 | 20 | 23039 | MULTISPECIES: HAD family |
| 10 | cont132 | 179.72 | 27 | 3.28E+04 | 19 | 19 | 66018 | spP04264 K2C1_HUMAN K |
| 10 | cont135 | 179.72 | 27 | 3.28E+04 | 19 | 19 | 66039 | crahCP1609934.2 keratin |
| 10 | cont134 | 179.72 | 27 | 3.28E+04 | 19 | 19 | 66067 | rfNP_006112.2 keratin 1 [H |
| 8 | WP_179004935.1 | 175.33 | 55 | 7.44E+04 | 17 | 17 | 21788 | MULTISPECIES: hypothetic |
| 11 | WP_179002106.1 | 155.45 | 43 | 3.96E+04 | 10 | 10 | 21441 | MULTISPECIES: chalcone i |
| 14 | WP_179002383.1 | 153.7 | 30 | 2.33E+04 | 8 | 8 | 19268 | MULTISPECIES: hypothetic |
| 14 | tr\|A0A2S1YTY4\|A0A2S1YTY4_9FLAO | 153.7 | 30 | 2.33E+04 | 8 | 8 | 19255 | Uncharacterized protein O |
| 14 | tr\|A0A1I1NVB8\|A0A1I1NVB8_9FLAO | 153.7 | 26 | 2.33E+04 | 8 | 8 | 23428 | Uncharacterized protein O |
| 12 | cont129 | 146.34 | 16 | 1.16E+04 | 10 | 10 | 58827 | trmQ8N175 Keratin 10 [Ho |
| 12 | cont133 | 146.34 | 16 | 1.16E+04 | 10 | 10 | 59528 | pirKRHU0 keratin 10 type I |
| 12 | cont122 | 146.34 | 16 | 1.16E+04 | 10 | 10 | 59519 | sptP13645 Keratin type I c |
| 12 | cont136 | 146.34 | 15 | 1.16E+04 | 10 | 10 | 63346 | crahCP1812051 keratin 10 |
| 20 | tr\|A0A2S1YFF8\|A0A2S1YFF8_9FLAO | 143.96 | 16 | 2.66E+03 | 6 | 6 | 36544 | Uncharacterized protein O |
| 20 | WP_179006530.1 | 143.96 | 16 | 2.66E+03 | 6 | 6 | 36507 | MULTISPECIES: hypothetic |
| 15 | WP_179003529.1 | 141.57 | 27 | 1.63E+04 | 8 | 8 | 33468 | MULTISPECIES: DUF4465 d |
| 19 | WP_179001695.1 | 140.13 | 12 | 6.44E+03 | 5 | 5 | 61467 | MULTISPECIES: tetratricop |
| 19 | tr\|A0A1I1S5A6\|A0A1I1S5A6_9FLAO | 140.13 | 12 | 6.44E+03 | 5 | 5 | 61457 | Tetratricopeptide repeat-c |
| 16 | WP_179008058.1 | 134.7 | 23 | 6.86E+03 | 7 | 7 | 33476 | MULTISPECIES: PKD domai |
| 16 | tr\|A0A1I1VPN8\|A0A1I1VPN8_9FLAO | 134.7 | 23 | 6.86E+03 | 7 | 7 | 33439 | PKD domain-containing pr |
| 16 | tr\|A0A2S1YFK7\|A0A2S1YFK7_9FLAO | 134.7 | 23 | 6.86E+03 | 7 | 7 | 33474 | PKD domain-containing pr |
| 22 | WP_179004570.1 | 114.38 | 12 | 9.05E+03 | 5 | 5 | 50462 | MULTISPECIES: DUF4302 d |
| 18 | WP_179004868.1 | 113.65 | 21 | 4.79E+03 | 5 | 5 | 25450 | MULTISPECIES: hypothetic |
| 17 | WP_179005968.1 | 112.25 | 23 | 6.33E+03 | 7 | 7 | 38984 | MULTISPECIES: SusE doma |
| 23 | ATCG00490.1 | 108.12 | 6 | 1.97E+04 | 3 | 3 | 52955 | large subunit of RUBISCO. |
| 21 | WP_179007264.1 | 94.27 | 22 | 2.14E+03 | 5 | 5 | 25962 | MULTISPECIES: DUF4197 d |
| 30 | ATCG00280.1 | 90.61 | 5 | 8.22E+02 | 2 | 2 | 51868 | chloroplast gene encoding |
| 26 | WP_179007155.1 | 83.32 | 16 | 3.03E+03 | 3 | 3 | 18998 | MULTISPECIES: DUF6265 f |
| 29 | CAA9197074.1 | 81.82 | 16 | 3.71E+03 | 2 | 2 | 22373 | Superoxide dismutase [Mn |
| 29 | WP_217427692.1 | 81.82 | 16 | 3.71E+03 | 2 | 2 | 22373 | superoxide dismutase [Fla |
| 29 | WP_179000806.1 | 81.82 | 16 | 3.71E+03 | 2 | 2 | 22392 | MULTISPECIES: superoxide |
| 29 | tr\|A0A1I1KNQ5\|A0A1I1KNQ5_9FLAO | 81.82 | 16 | 3.71E+03 | 2 | 2 | 22429 | Superoxide dismutase OS= |
| 29 | tr\|A0A2S1YI81\|A0A2S1YI81_9FLAO | 81.82 | 16 | 3.71E+03 | 2 | 2 | 22429 | Superoxide dismutase OS= |
| 27 | AT2G21330.2 | 72.8 | 7 | 9.90E+02 | 2 | 2 | 33323 | fructose-bisphosphate ald |
| 27 | AT2G21330.3 | 72.8 | 6 | 9.90E+02 | 2 | 2 | 41808 | fructose-bisphosphate ald |
| 27 | AT2G21330.1 | 72.8 | 6 | 9.90E+02 | 2 | 2 | 42931 | fructose-bisphosphate ald |
| 24 | tr\|A0A2S1YIE6\|A0A2S1YIE6_9FLAO | 72.26 | 10 | 7.17E+02 | 3 | 3 | 33197 | LD-carboxypeptidase OS= |
| 24 | WP_179000684.1 | 72.26 | 10 | 7.17E+02 | 3 | 3 | 33096 | MULTISPECIES: LD-carbox |
| 24 | tr\|A0A1I1KK42\|A0A1I1KK42_9FLAO | 72.26 | 9 | 7.17E+02 | 3 | 3 | 37036 | Muramoyltetrapeptide car |
| 25 | WP_179005419.1 | 69.98 | 21 | 3.30E+03 | 3 | 3 | 19411 | MULTISPECIES: DinB family |
| 34 | AT1G07930.2 | 61.32 | 5 | 1.19E+03 | 2 | 2 | 41374 | elongation factor 1-alpha / |
| 34 | AT5G60390.2 | 61.32 | 5 | 1.19E+03 | 2 | 2 | 44356 | elongation factor 1-alpha / |
| 34 | AT1G07940.1 | 61.32 | 4 | 1.19E+03 | 2 | 2 | 49502 | elongation factor 1-alpha / |
| 34 | AT1G07930.1 | 61.32 | 4 | 1.19E+03 | 2 | 2 | 49502 | elongation factor 1-alpha / |
| 34 | AT5G60390.3 | 61.32 | 4 | 1.19E+03 | 2 | 2 | 49502 | elongation factor 1-alpha / |
| 34 | AT5G60390.1 | 61.32 | 4 | 1.19E+03 | 2 | 2 | 49502 | elongation factor 1-alpha / |
| 34 | AT1G07940.2 | 61.32 | 4 | 1.19E+03 | 2 | 2 | 49502 | elongation factor 1-alpha / |
| 34 | AT1G07920.1 | 61.32 | 4 | 1.19E+03 | 2 | 2 | 49502 | elongation factor 1-alpha / |
| 33 | tr\|A0A2S1YL73\|A0A2S1YL73_9FLAO | 60.81 | 6 | 2.03E+03 | 2 | 2 | 43065 | Elongation factor Tu OS=Fl |
| 33 | WP_026730432.1 | 60.81 | 6 | 2.03E+03 | 2 | 2 | 43008 | MULTISPECIES: elongation |
| 33 | tr\|A0A1I1PYR9\|A0A1I1PYR9_9FLAO | 60.81 | 6 | 2.03E+03 | 2 | 2 | 43083 | Elongation factor Tu OS=Fl |
| 31 | cont127 | 56.74 | 4 | 1.06E+03 | 2 | 2 | 60092 | sptP48668 Keratin type II c |
| 31 | cont125 | 56.74 | 4 | 1.06E+03 | 2 | 2 | 59914 | sptP02538 Keratin type II c |
| 31 | cont126 | 56.74 | 4 | 1.06E+03 | 2 | 2 | 60069 | sptP48666 Keratin type II c |
| 36 | AT1G67090.1 | 53.83 | 8 | 1.08E+04 | 2 | 2 | 20216 | RBCS1A (RIBULOSE BISPH |
| 32 | ATCG00120.1 | 53.59 | 4 | 6.78E+02 | 2 | 2 | 55328 | Encodes the ATPase alpha |

**Supplemental Table 1.** Identity of constituent proteins within the ~21 kDa band separated from Nagase crude alginate lyse preparation.

### Supplemental Table 2

| **Ligand full name** | **Ligand short name** | ***Fq*PBP** | **HusA** |
|---|---|---|---|
| Coproporphyrinogen III | **cgen3** | -8.4702 | -7.6545 |
| Protoporphyrinogen IX | **pgen9** | -8.9597 | -8.0984 |
| Protoporphyrin IX | **ppor9** | -10.4674 | -7.8561 |
| Heme B | **hemeb** | -10.1500 | -8.1855 |
| Hemin chloride (Heme-Fe3+, Cl-) | **hem3** | -9.3278 | -8.5247 |
| Hemin no Cl- (oxidized Heme B) | **hem3x** | -10.2383 | -8.3944 |
| Zinc Protoporphyrin | **porzo** | -9.2250 | -8.3916 |
| Zinc Protoporphyrin-H2O | **porzn** | -10.5126 | -8.5455 |
| Chlorophyll A | **chloa** | -9.553 | -7.1747 |
| Chlorophyll B | **chlob** | -9.9153 | -6.6579 |

**Supplemental Table 2.** Docking scores (kcal/mol) of *Fq*PBP and HusA against porphyrin related ligands obtained from AutoDock Vina (More negative values indicate higher binding affinity).

### Supplemental Figure 1

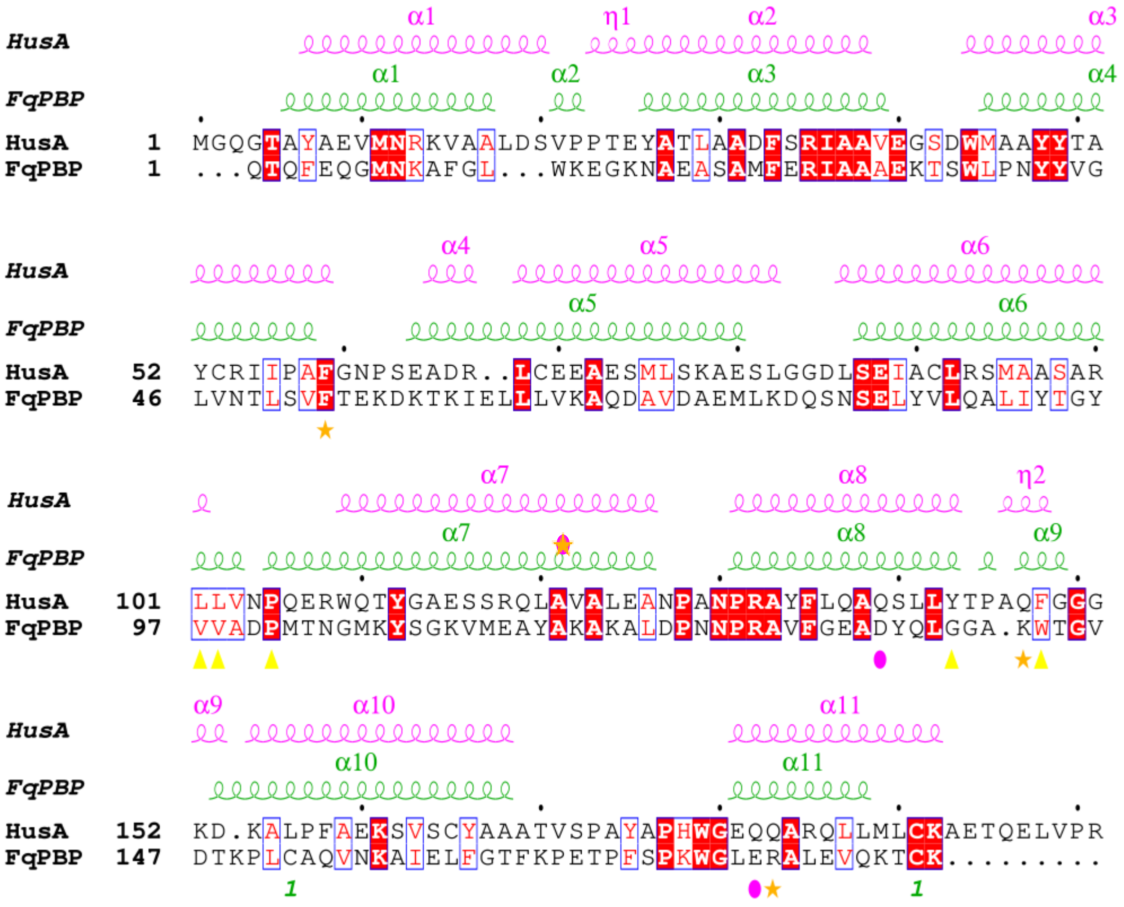


**Supplemental Figure 1.** Sequence alignment of *Fq*PBP and HusA generated with ClustalO including the secondary structures. The major binding contributing residues found by MMPBSA with Protoporphyrin IX on *Fq*PBP are shown below the sequence by the symbols ★, ▲ and ●. The marks *1* indicate a cysteine bridge in *Fq*PBP.
